# An intragastric murine model that recapitulates natural infection with invasive non-typhoidal *Salmonella* for preclinical evaluation of vaccine immunogenicity and protection

**DOI:** 10.64898/2026.11.22.768488

**Authors:** Valentina Mocci, Fabio Fiorino, Martina Tambassi, Giulia D’annunzio, Laura Mazzera, Giorgio Montesi, Andrea Luppi, Stefano Pongolini, Susanna Ricci, Gianni Pozzi, Donata Medaglini, Elena Pettini

## Abstract

Invasive non-typhoidal *Salmonella* disease is a major global health concern, particularly in sub-Saharan Africa, where young children and immunocompromised individuals are particularly vulnerable. Preclinical models that reproduce the natural course of invasive non-typhoidal *Salmonella* infection without antibiotic-mediated disruption of the intestinal microbiota are essential for vaccine development. Here, an antibiotic-free murine model is described based on intragastric infection of susceptible C57BL/6 mice with the ST313 clinical isolate *Salmonella enterica* serovar Typhimurium D23580. Over seven days, infection progressed from intestinal colonization to extraintestinal dissemination, accompanied by increasing histopathological damage and activation of local and systemic inflammatory responses. The ability of the infection model to evaluate vaccine-induced immunity and protection was assessed by immunizing mice with a formalin-inactivated D23580 strain. Vaccination elicited *Salmonella*-specific serum IgG and reduced clinical severity, with vaccinated mice showing significantly lower bacterial loads in blood, liver and colon compared with control mice after intragastric challenge.

By combining the enteric route, which mimics the natural route of human infection, with an intact intestinal microbiota, this model reproduces key features of progressive invasive non-typhoidal *Salmonella* disease and provides a tool for investigating host–pathogen interactions and evaluating vaccine-induced immunity and protection.

**Importance:** *Salmonella enterica* serovar Typhimurium commonly causes diarrhoeal illness, but in sub-Saharan Africa it can cause life-threatening bloodstream infections, particularly in young children and immunocompromised people. No vaccine is licensed for this disease. Animal models are needed to study infection progression and evaluate vaccine induced immunity and protection under relevant conditions. Many mouse models use antibiotic pretreatment that alters the intestinal microbiota. In this model, bacteria are administered intragastrically in the absence of antibiotics, thereby recapitulating the gastrointestinal phase of infection while avoiding deliberate disruption of the gut microbial community. Infection progressed from the intestine to organs beyond the intestine, with tissue damage and inflammation consistent with invasive disease. A prototype vaccine was used as a proof-of-concept intervention and reduced disease severity and bacterial spread. The model provides a tool that faithfully recapitulates the gastrointestinal phase of invasive *Salmonella* disease and supports evaluation of vaccine-induced protection and other interventions.

## INTRODUCTION

*Salmonella* species are facultative intracellular Gram-negative bacteria that cause different types of disease in different hosts. *Salmonella enterica* subspecies *enterica* serovar Typhi (*S*. Typhi) is a strict human pathogen responsible for typhoid fever, an invasive life-threatening infection that mainly affects individuals ranging from infants to young adults (1). In 2021, *S.* Typhi caused approximately 7.2 million cases of enteric fever and 93,000 deaths worldwide (2). By contrast, non-typhoidal *Salmonellae* (NTS) comprise a broad group of *Salmonella* serovars other than the typhoidal serovars, including *Salmonella enterica* serovars Typhimurium and Enteritidis, which are among the most common causes of human NTS disease (3, 4). In immunocompetent individuals, NTS infections most frequently present as self-limiting diarrhoea and pathologically characterized by enterocolitis, mainly involving the distal ileum and colon (5). In immunocompromised subjects, however, NTS are often associated with invasive infection that often presents as septicaemia, anaemia, kidney damage and encephalopathy with no or limited enteric symptoms (5). Co-infections (HIV, malaria), immunosuppression and other conditions (malnutrition, anaemia) are considered as risk factors for invasive non-typhoidal *Salmonella* disease (iNTS) (3). The burden of iNTS disease was estimated to be 510,000 cases and 62,000 deaths in 2021 (6), with incidence being particularly high and geographically heterogeneous across sub-Saharan Africa (7, 8). Phylogenetic analysis has shown that *S*. Typhimurium isolates associated with iNTS disease predominantly belong to sequence type 313 (ST313), a lineage of *S*. Typhimurium widely distributed in sub-Saharan Africa (9, 10). Despite growing global efforts to develop effective vaccines against iNTS disease, no vaccine is currently licensed for human use. However, several vaccine candidates based on different technological platforms, including glycoconjugate and GMMA-based vaccines, are currently undergoing early-phase clinical evaluation, with GMMA formulations showing promising safety and immunogenicity profiles in first-in-human studies (11, 12).

In this context, preclinical murine models remain essential for evaluating the immunogenicity and protective efficacy of vaccine candidates against invasive *Salmonella* infection, providing a critical tool for investigating disease pathogenesis and assessing potential therapeutic and prophylactic interventions (13, 14). These infection models vary depending on whether they are intended to reproduce typhoid-like systemic disease or gastroenteritis/enterocolitis (15, 16). In mice, systemic typhoid-like fever is generally reproduced using a surrogate disease model in which either genetically susceptible (e.g., Nramp1 deficient strains, such as C57BL/6, BALB/c, DBA/1) or resistant (e.g., CBA, 129Sv, C3H/Hen, DAB/2) mouse strains are infected with either *S.* Typhimurium or *S.* Enteritidis. These infections result in a systemic disease characterized by limited intestinal inflammation and progressive extraintestinal dissemination, resembling features of human typhoid fever (17–20). While both susceptible and resistant mouse strains can exhibit systemic dissemination, resistant strains more effectively control bacterial replication thereby limiting disease severity and often resulting in a persistent infection state (16, 21).

Conversely, enterocolitis models are based on disrupting the intestinal microbiota, typically through antibiotic pretreatment with streptomycin or the use of germ-free mice, followed by oral infection with *S.* Typhimurium (22–25). In these models, inflammation is predominantly localized in the cecum and colon, in contrast to the small intestine being the most affected site in human gastroenteritis (26, 27). Additionally, streptomycin-pretreated mice can develop chronic colitis, cholangitis, and bacterial persistence in the gallbladder, resembling the chronic carrier state of *S.* Typhi in humans (25, 28, 29). All current mouse infection models face limitations in fully replicating the conditions of natural infection. Many models require oral pretreatment with antibiotic and/or sodium bicarbonate to reduce the intestinal microbiota and/or to neutralize gastric acidity, respectively (22–25, 30). Furthermore, some models bypass the intestinal mucosa by using parenteral infection routes (e.g., intraperitoneal, intravenous, or subcutaneous), which do not reflect the natural route of infection (31, 32).

The present study describes the development and characterization of a murine model of iNTS disease in susceptible C57BL/6 mice based on intragastric infection with the ST313 clinical isolate *S*. Typhimurium D23580, without prior streptomycin pretreatment. This strain, belonging to the ST313 clade, is associated with adaptation to invasive disease in immunocompromised subjects (9, 10). D23580 displays genomic and phenotypic features linked to enhanced systemic infection and is characterized by an increased ability to cause bacteraemia and to colonize systemic sites following intragastric infection, closely resembling features of iNTS disease in vulnerable human populations (33).

To characterise the acute course of infection and assess the utility of the model, clinical disease, bacterial dissemination, tissue pathology, local inflammatory gene expression, and serum cytokine and chemokine responses were monitored at predefined time points.

The suitability of the model as a tool for the preclinical evaluation of interventions was subsequently assessed in two vaccine-associated cohorts, using vaccination as a proof-of-concept application. In the first cohort, a prototype formalin-inactivated D23580 preparation was evaluated for its ability to induce antigen-specific antibody responses and to reduce disease severity and bacterial dissemination following challenge. This experiment was intended to test the suitability of the model for detecting intervention-associated effects, rather than to provide a comprehensive evaluation of the preparation as a vaccine candidate. In a separate long-term cohort, the immunological characterization was extended by analysing selected B-cell populations in the spleen and draining lymph nodes six months after priming and six days after D23580 challenge.

## MATERIAL AND METHODS

### Bacterial strains and culture conditions

The invasive clinical isolate from Malawi *S*. Typhimurium D23580 (9, 34) was grown in Luria-Bertani (LB) broth [tryptone (10 g/L, Oxoid, Basingstoke, UK), sodium chloride (10 g/L, Carlo Erba, Milan, Italy), yeast extract (5 g/L, Oxoid)] up to optical density at 590nm (OD)_590 =_ 0.35, and then stored at -80 °C with 10% glycerol (Carlo Erba) until use, as previously described (13). Colony forming units (CFU) were determined by plating serial dilutions onto LB agar plates and incubating overnight at 37 °C with 5% CO_2._ In infection experiments, chloramphenicol (64 µg/mL; Sigma-Aldrich) was added to LB agar plates to selectively quantify *S.* Typhimurium D23580 recovered from cecum and colon samples and simultaneously suppress commensal bacterial growth, whereas samples from blood, spleen, and liver were plated on non-selective LB agar. The chloramphenicol concentration was determined by preliminary minimum inhibitory concentration (MIC) assays (data not shown).

### Animal cohorts and experimental design

Eight-week-old female C57BL/6 (Nramp1⁻/⁻) mice were purchased from Charles River Italy (Lecco, Italy) and maintained under pathogen-free conditions in the animal facility of the Laboratory of Molecular Microbiology and Biotechnology (LA.M.M.B.), Department of Medical Biotechnologies, University of Siena (Italy). Animals were housed in groups of up to 8 per cage (Tecniplast, Buguggiate, Italy) at 20–24 °C and 55 ± 10% humidity, with food and water available ad libitum, under conditions complying with the “3R” principles in accordance with the Italian national guidelines (D. Lgs. no. 26/2014) and in conformity with European Directive 2010/63/EU on the protection of animals used for scientific purposes. The study was approved by the Italian Ministry of Health (authorization no. 371/2019-PR).

A total of 108 female C57BL/6 mice were included in two main experimental cohorts (Figure 1), comprising 104 experimental mice and four additional naïve reference control mice. The first cohort (Cohort 1) comprised 94 mice and was divided into three experimental arms according to the primary purpose of the experiment named Cohort 1A, Cohort 1B and Cohort 1C respectively. Cohort 1A, the infection-model characterization arm, included 24 mice that were intragastrically infected with *S.* Typhimurium D23580 and euthanized at 1, 3 or 7 days post-infection. Four additional naïve, uninfected mice were included in Cohort 1A as a reference control for histopathological and tissue gene-expression analyses. The 24 infected mice in Cohort 1A were used to characterize the clinical progression of infection, bacterial dissemination to intestinal and systemic organs, tissue histopathology, local inflammatory gene expression and serum cytokine and chemokine profiles. Cohort 1B, the cytokine-resolution arm, included an additional 18 infected mice maintained under the same experimental conditions and euthanized at 3, 5 or 7 days post-infection. This arm was included to provide additional temporal resolution for the characterization of circulating cytokine and chemokine kinetics.

**Figure 1.**
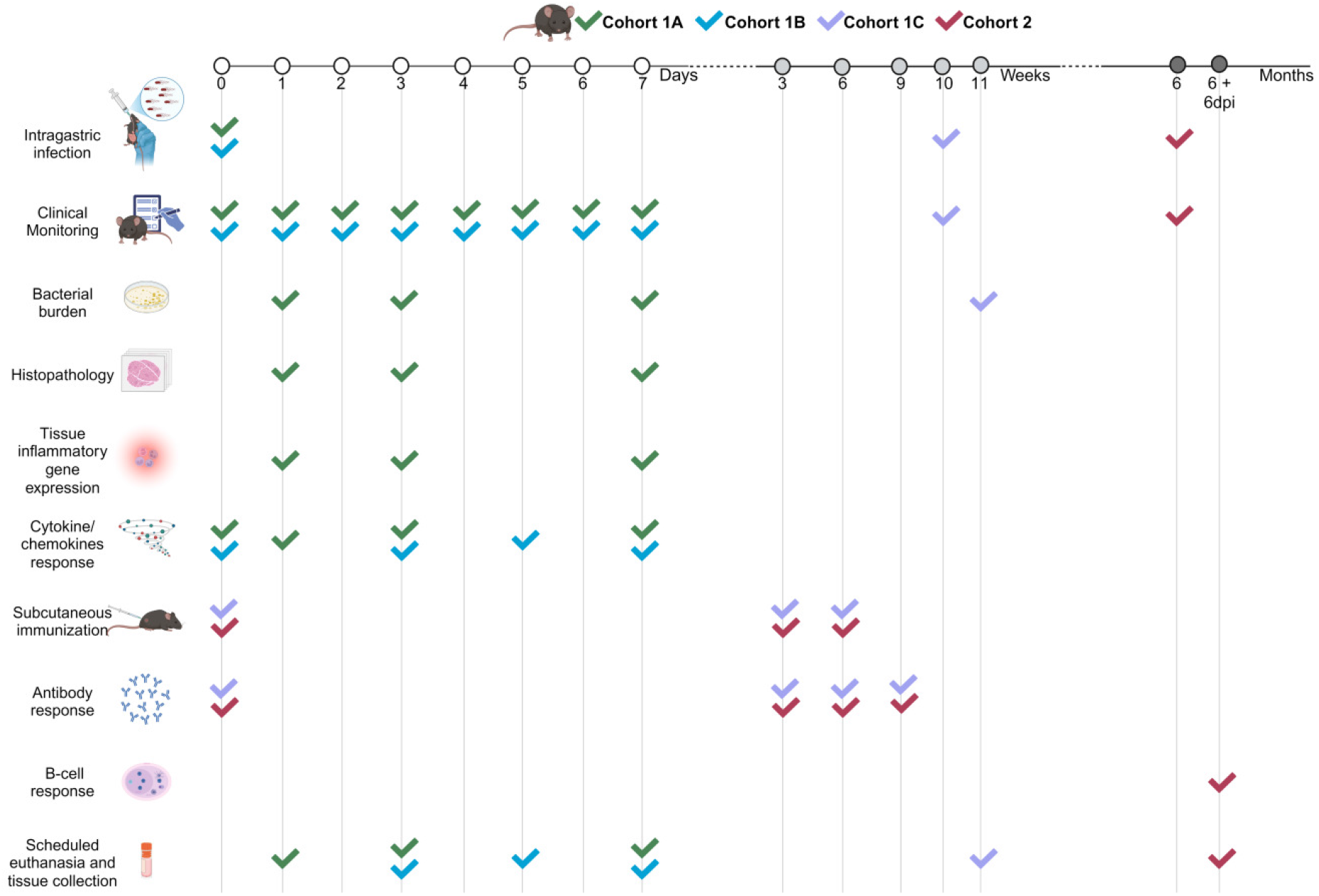
Experimental workflow and animal cohorts. The study comprised two main cohorts, identified by distinct colours in the schematic. Cohort 1 included three subcohorts. Cohort 1A (green; n = 24) was used to characterize infection progression after intragastric challenge with live *Salmonella* Typhimurium D23580, with scheduled euthanasia at 1, 3 and 7 days post-infection (dpi). Cohort 1B (blue; n = 18) was euthanized at 3, 5 and 7 dpi to provide additional temporal resolution for serum cytokine and chemokine profiling. Cohort 1C (purple; n = 48) was used as a proof-of-concept intervention cohort to evaluate a formalin-inactivated D23580 vaccine preparation, including longitudinal measurement of serum anti-*Salmonella* IgG and assessment of clinical and microbiological outcomes after challenge. A subset of vaccinated mice (n = 8) and saline-treated controls (n = 8) was challenged with live D23580 ten weeks after the first immunization and euthanized 6 dpi. Cohort 2 (red; n = 14) was used to assess long-term vaccine-associated B-cell responses. Vaccinated mice (n = 8) and saline-treated controls (n = 6) followed the same immunization schedule; serum IgG levels were measured at weeks 0, 3, 6 and 9, and mice were challenged six months after the first immunization and euthanized 6 dpi (6 + 6dpi) for multiparameter flow-cytometric analysis of the spleen and draining lymph nodes. Check marks indicate the cohorts contributing to each experimental procedure. Open circles indicate days, light-grey circles indicate weeks, and dark-grey circles indicate months; these units refer to days post-infection, weeks after the first immunization and months after the first immunization, respectively. Four additional naïve, uninfected mice served as day-0 reference controls for the histopathological and tissue gene-expression analyses associated with Cohort 1A. These animals were not included in the n = 24 reported for Cohort 1A and are not shown in the schematic. Created with BioRender.com.

The third arm, Cohort 1C, was used for vaccine assessment, it included 48 mice and was used to evaluate the immunogenicity and protective efficacy of a prototype formalin-inactivated *S.* Typhimurium D23580 vaccine. Vaccinated and saline-treated control animals were immunized according to the vaccination schedule described below. A defined subset of vaccinated (n = 8) and control (n = 8) animals was subsequently challenged intragastrically with live *S.* Typhimurium D23580 and monitored for clinical disease, bacterial dissemination and IgG production.

The second cohort, Cohort 2, comprised 14 additional mice and was used to assess the long-term B-cell response induced by vaccination. This cohort included eight vaccinated mice and six saline-treated control mice. Six months after the first immunization, vaccinated and control animals were challenged intragastrically with live *S.* Typhimurium D23580 using the same infection model applied in Cohort 1. All animals in Cohort 2 were euthanized six days after infection, and the spleen and draining lymph nodes were collected for multiparameter flow-cytometric analysis of B-cell responses.

### Intragastric infection by oral gavage

Mice in Cohorts 1A and 1B were intragastrically infected by oral gavage on day 0 with 10^8 CFU per mouse of *S.* Typhimurium D23580. Mice in Cohorts 1C and 2 were infected after completion of the corresponding immunization schedule, at the time points indicated in Figure 1. The infectious dose was selected on the basis of previous studies conducted in our laboratory (Supplementary Figure 1) and was consistent with previous murine infection studies using the D23580 strain (33). Bacteria were administered in a total volume of 100 μl/mouse in LB broth solution by using a single-use gavage needle (20 GA, 38 mm length, 1.75 mm tip diameter) as previously described (35). The inoculum size was verified by plating on LB agar plates.

### Clinical monitoring and body weight assessment

Following intragastric infection, mice from Cohorts 1A, 1B, 1C and 2 were monitored daily for body weight changes and clinical signs of disease. Body weight variation was calculated for each animal as the percentage change relative to the body weight recorded before infection. Clinical signs were assessed using a score ranging from 0 to 5, adapted from previously described criteria (36). The score was assigned based on the evaluation of different markers, including fur appearance, behaviour, activity, posture, eyelid closure and righting reflex. Specifically, disease severity was classified as follows: 0, absence of clinical signs; 1, piloerection and reduced motility; 2, hunched posture and semi-closed eyes; 3, delayed righting reflex, defined as more than 5 seconds to return from the supine to the prone position; 4, inability to return from the supine to the prone position; and 5, clinical signs requiring euthanasia according to the predefined humane endpoint criteria. Humane endpoint criteria were predefined in the approved animal protocol. In accordance with these criteria, mice were humanely euthanized when any of the following conditions was reached: severe clinical deterioration corresponding to a clinical score of 5, persistent clinical score of 4 on two consecutive daily evaluations, or body-weight loss ≥25%. Deaths occurring before the next scheduled monitoring time point were recorded as mortality events.

### Analysis of bacteria dissemination to intestinal and peripheral tissues

Mice were euthanized at predefined time points after infection by intraperitoneal administration of an overdose of Ketamine (Lobotor, 100 mg/kg; ACME srl, Italy,) and Xylazine (Rompun, 8 mg/kg; Bayer SPA, Germany). For bacterial dissemination analyses, animals from Cohort 1A were euthanized at 1, 3 or 7 days post-infection, whereas the challenged subset of Cohort 1C was euthanized at 6 days post-infection (Figure 1). Cohort 1B was euthanized at 3, 5 or 7 days post-infection for serum cytokine and chemokine profiling, while Cohort 2 was euthanized 6 days after challenge for long-term B-cell analyses. Blood samples were collected by cardiac puncture using a 1 ml syringe 25GA x 5/8in (BD PlastipakTM, Becton Dickinson USA) and stored at -80°C in LB broth with 10% glycerol. At different time points, liver, spleen and the intestine (further sectioned into ileum, cecum and colon) were aseptically removed with veterinary forceps and scissors and divided into predefined portions for CFU counts and histological analysis. The organs were homogenized in LB broth + 10% glycerol by Ultra-Turrax® T-10 basic homogeniser (IKA®-Werke, Staufen, Germany) at 28800 rpm and stored at -80 °C in LB broth with 10% glycerol. Bacterial loads were determined by plating 10-fold serial dilutions onto LB agar plates and incubating at 37 °C for 24 h, using an approach adapted from previously described methods for quantifying *Salmonella* burden in murine organs (37). Cecum, colon, spleen and liver were weighed prior to homogenization.

### Histological Analysis

Tissue portions of cecum, liver, and spleen from mice in Cohort 1A were fixed in 10% neutral buffered formalin, embedded in paraffin blocks, and sectioned in 3 μm slices. The resulting sections were mounted on glass slides and stained with hematoxylin and eosin (H&E) according to standard procedures. Blinded evaluation of the stained tissue sections was performed independently by two pathologists. A semi-quantitative analysis of cecal lesions was conducted using a scoring system (ranging from 0 to 3) modified from the literature (23, 30). The parameters evaluated included the submucosal edema, inflammation, goblet cell number, and epithelial integrity (Supplementary Table S1). For each cecal sample, the scores of each parameter were summed, and the final pathological score of each sample was calculated as the average of the two independent scores, ranging from 0 to 12, where: 0 = no inflammation; 1 to 2 = minimal signs of inflammation; 3 to 4 = slight inflammation; 5 to 8 = moderate inflammation; 9 to 12 = profound inflammation. Hepatic and splenic inflammation and necrosis were evaluated as previously reported (37) (Supplementary Table S2), and their pathological score sample was the mean of the sum of two independent pathological evaluations. Additionally, the liver and spleen were examined for the presence of vascular thrombi, as thrombosis is a recognized complication of systemic infections (38). Four additional naïve, uninfected mice were processed identically and served as reference controls for the histopathological and tissue gene-expression analyses.

### Tissue cytokine gene-expression analysis

Portions of cecum and liver collected from mice in Cohort 1A were submerged in RNAlater (Qiagen) and stored at -20°C until use. Cecum and liver samples from the same four naïve reference mice were processed identically and used as the reference group for tissue gene-expression analyses. Total RNA was extracted using MagMAX™ *mir*Vana™ Total RNA Isolation Kit (Applied Biosystems) and the KingFisher™ Apex Purification System with 96 Deep–Well Head (Thermo Scientific). RNA concentration and quality (A_260/_A_280)_ were determined through a Synergy H1 Hybrid spectrophotometer (BioTek) equipped with Gen5 software. Reverse transcription was performed by using the High-Capacity cDNA Reverse Transcription Kit (Applied Biosystems). Final cDNA was diluted 1:10 for PCR reactions. Quantitative real-time PCR (qRT-PCR) was performed using GoTaq qPCR Master Mix (Promega Corporation) and the Biorad CFX-96 thermocycler. All reactions were run in duplicate. The following primers sequences (5′-3′) were used: GAPDH forward TGTAGACCATGTAGTTGAGGTCA; GAPDH reverse AGGTCGGTGTGAACGGATTTG; KC forward ATGGCTGGGATTCACCTCAA; KC reverse AGTGTGGCTATGACTTCGGTTT; TNF forward CCACCACGCTCTTCTGTCTAC; TNF reverse AGGGTCTGGGCCATAGAACT. Transcription of the target genes (KC and TNF) was normalized to the reference gene coding for the glyceraldehyde-3-phosphate dehydrogenase (GAPDH) (39, 40). The Common Base Method was used for the analysis of qRT-PCR data (41).

### Multiplex Cytokine and Chemokine Assay

Serum concentrations of IL-1α, IL-1β, IL-2, IL-3, IL-4, IL-5, IL-6, IL-9, IL-10, IL12(p40), IL-12(p70), IL-13, IL17A, eotaxin, G-CSF, GM-CSF, IFN-γ, KC, MCP-1, MIP-1α, MIP-1β, RANTES and TNF-α were assessed using a Luminex immunoassay. Blood samples were collected from mice in Cohorts 1A and 1B before infection and at the scheduled experimental endpoint. Pre-infection samples were assigned to day 0, while data from the Cohorts 1A and 1B were combined at the shared time points to characterize systemic cytokine and chemokine dynamics at days 0, 1, 3, 5, and 7 post-infection. Blood samples were centrifuged at 1200× *g* at 4 °C for 15 min, and sera were collected and stored at -80 °C. Analytes were detected using the BioPlex pro mouse cytokine group 1-panel 23-plex immunoassay (Bio-Rad) following the manufacturer’s protocol and analysed by the Bio-Plex 200 Reader (Bio-Rad). Cyto- and chemokine concentrations, expressed as pg/mL, were calculated based on standard curves using Bio-Plex Manager 6.2.

An exploratory multivariate analysis, integrating serum cytokine and chemokine profiles from Cohorts 1A and 1B with surrogate mortality-risk labels derived from a previously developed model (42), was carried out and is detailed in the Supplementary Methods and Supplementary Results.

### *S.* Typhimurium inactivation and animal immunization

Whole-cell killed vaccines represent a long-established and widely used approach in vaccinology. Among these, preparations based on chemically inactivated *Salmonella*, including formalin-inactivated whole-cell vaccines, have previously been used in murine models to assess vaccine-induced immune responses and protection (32, 43, 44). In the present study, the vaccine preparation protocol was adapted with minor modifications from previously published procedures and described below. *S.* Typhimurium D23580 was cultured in LB broth at 37 °C with agitation (95 rpm) until OD_590 =_ 0.4. The culture was centrifuged (6000 × g, 30 min, 4 °C), washed with sterile distilled water, and centrifuged again. The pellet was resuspended in sterile distilled water, and CFU/ml was determined. Bacteria were inactivated by incubation with 2% formaldehyde (Carlo Erba), followed by overnight incubation (37°C, 300 rpm). The sample was centrifuged (6000 × g, 20 min, 4 °C), washed with sterile distilled water, resuspended in sterile distilled water (Fresenius Kabi, Germany) and stored at -80 °C until use.

To confirm bacterial inactivation, aliquots of the vaccine formulation were plated onto LB. Mice from Cohorts 1C and 2 were immunized subcutaneously with 10^8 CFU-equivalent/mouse in 100 µL of the inactivated *S*. Typhimurium D23580 strain at weeks 0, 3, and 6. Control mice received 100 µL of saline.

### ELISA assay

Maxisorp microtiter 96-well plates (Nunc, Roskilde, Denmark) were coated with 5 x 10^6 CFU equivalent/ml of the inactivated *S*. Typhimurium D23580 strain in a carbonate buffer pH 9.4 (Sigma-Aldrich, St. Louis, MO, US), and incubated overnight at 4 °C. Coating was removed, and plates were blocked with 200 µL/well of PBS containing 0.05% Tween 20 (Sigma-Aldrich) and 5% fat-free milk (AppliChem, Darmstadt, Germany) for 2 h at 37 °C. After washing with PBS-Tween 20, serum samples collected from mice in Cohorts 1C and 2 at weeks 0, 3, 6 and 9 were added and titrated in twofold dilutions in duplicate in PBS containing0.05% Tween 20 and 2.5% fat-free milk (diluentbuffer) in 100 µL/well. After incubation for 1 h at 37 °C, plates were washed, incubated for 1 h at 37 °C with alkaline phosphatase-conjugated goat anti-mouse IgG (diluted 1:1500; Southern Biotechnology, Birmingham, AL, USA) in 100 µL/well and developed by adding 1 mg/mL of alkaline phosphatase substrate (Sigma-Aldrich) in 200 µL/well. The optical density was measured at 405 nm and recorded using a Multiskan FC Microplate Photometer (Thermo Scientific, Waltham, MA, USA). Antibody end point titres were reported as the reciprocal of the highest sample dilution with an OD value at least twofold higher than the background.

### Multiparametric Flow Cytometric Analysis

Following euthanasia, the spleen and draining lymph nodes were harvested from mice in Cohort 2, manually disaggregated and filtered through nylon mesh filters to obtain single-cell suspensions. Cells from spleen and draining lymph nodes were used to characterize B-cell associated responses induced by immunization and challenge. Samples were incubated for 30 min at 4° C in Fc-blocking solution [CD16/CD32 mAb (clone 93; eBioscience) 1:1000 diluted in cRPMI]. Cells were then washed in cRPMI and then labelled with live/dead Fixable Viability Stain 780 according to the manufacturer’s instruction (1:2000 diluted in PBS; BD Biosciences). Cells were subsequently washed and surface stained with different combination of antibodies: PE-CF594-conjugated anti-CD3 (clone 145-2C11), BUV395-conjugated anti-B220 (clone RA3-6B). AF647-conjugated anti-CD38 (90/CD38), FITC-conjugated anti-GL7 (clone GL7), R718-conjugated anti-CD95 (clone Jo2), BV421-conjugated TACI (clone 8F10), BV786-conjugated anti-CD138 (clone CD281-2), RB705-conjugated anti-CXCR4 (clone 2B11/CXCR4), BV711-conjugated anti-IgM (clone R6-60.2), BV605-conjugated anti-IgD (clone 11-26c.2a) and BV650-conjugated anti-IgG (clone A85-1), all from BD Biosciences (New York, NY, USA). Cells from different samples were washed twice in PBS, Fix Perm was then added for 10 min at 4° C, and cells were resuspended in 250 µl in filtered PBS. All antibodies were titrated before use to determine the optimal dilution.

Fluorescence-minus-one controls were used to detect the background fluorescence for each fluorochrome. Approximately 500.000 events were acquired and stored for each sample with SO LSRFortessa X20 flow cytometer (BD Biosciences). Data analysis was performed using FlowJo v10 (TreeStar, Ashland, OR, USA).

### Statistical analysis

Survival was analysed using the Kaplan–Meier method. Bacterial loads, expressed as CFU per organ or CFU/g of tissue or faecal content, are presented as mean ± standard error of the mean (SEM). For comparisons involving more than two independent groups, differences in bacterial burden across time points were assessed using the Kruskal-Wallis test followed by Dunn’s multiple-comparisons test. Direct comparisons between two independent groups, including vaccinated and saline-treated control mice, were performed using two-tailed Mann-Whitney U tests.

For each tissue, differences in histopathological scores among infected groups at different time points and naïve controls were assessed using the Kruskal-Wallis test followed by Dunn’s multiple-comparisons test. Fisher’s exact test was used to evaluate differences between groups in the dichotomous variable of vascular thrombosis presence/absence in the liver and spleen. Differences in gene expression were assessed using one-way ANOVA followed by Tukey’s multiple-comparisons test. Serum cytokine and chemokine concentrations were log10-transformed before analysis. For each mediator, temporal differences across days 0, 1, 3, 5 and 7 post-infection were assessed separately using one-way ANOVA followed by Tukey’s multiple-comparisons test. Serum antibody levels were expressed as geometric mean titers (GMT) with 95% confidence intervals, and differences across the vaccination time course were assessed using the Kruskal-Wallis test followed by Dunn’s multiple-comparisons test.

For flow-cytometry analyses, direct comparisons between vaccinated and control mice were performed using exact two-tailed Mann–Whitney U tests. All statistical analyses were performed using GraphPad Prism version 10.3.0 (GraphPad Software Inc., San Diego, US). For all analyses, statistical significance was defined as *P ≤ 0.05; **P ≤ 0.01; ***P ≤ 0.001; ****P ≤ 0.0001.

## RESULTS

The present study comprised two main study cohorts (Figure 1). Cohort 1 comprised 90 experimental mice and four additional naïve, uninfected reference mice (94 animals in total) and was subdivided into three experimental arms. Cohort 1A included 24 infected mice used to characterize clinical progression, bacterial dissemination, tissue pathology and local inflammatory gene expression after intragastric infection, with scheduled euthanasia at 1, 3 and 7 days post-infection. Cohort 1B (n = 18) comprised an additional infected group with scheduled euthanasia at 3, 5 and 7 days post-infection and was used to increase the temporal resolution of serum cytokine and chemokine profiling; serum measurements from Cohorts 1A and 1B were combined at shared time points. To provide a proof-of-concept application of the infection model, Cohort 1C (n = 48) was used to assess vaccine-associated changes in antibody responses, clinical disease and bacterial dissemination after challenge. Cohort 2 (n = 14; eight vaccinated and six saline-treated control mice) was used to characterize selected B-cell populations following the long-term vaccination and intragastric challenge with *S.* Typhimurium D23580.

### Intragastric D23580 infection induces progressive disease and systemic bacterial dissemination

Cohort 1A, the infection-model characterization arm, was used to define the clinical, microbiological and pathological progression of intragastric *S.* Typhimurium D23580 infection. Mice were monitored daily and euthanized at 1, 3 or 7 days post-infection, allowing bacterial dissemination, tissue pathology and local inflammatory responses to be assessed across the acute phase of infection.

Body weight changes and clinical scores (e.g., fur appearance, eyelids, activity, posture, behaviour) are considered valuable indicators of health status in infected animals (36, 45). Therefore, both readouts were daily monitored in mice upon enteric infection to analyse disease progression. Infected mice showed a progressive and sustained reduction in body weight over time (Figure 2A). Body weight loss was detectable from day 2 post-infection and reached up to 25% of the initial body weight by day 5. This coincided with the onset of mortality, as 2 out of 8 animals died at this time point (Figure 2A). Clinical scores closely mirrored weight loss dynamics, with marked health deterioration observed from day 5 post-infection (Figure 2B). In order to evaluate bacterial dissemination kinetics following infection, blood, liver, spleen, cecum and colon were collected at 1, 3 and 7 days after infection (Figure 2C-D). One day post-infection, no bacteria were detected in extraintestinal organs, whereas early gut colonization was observed in both the cecum and colon of 63% (1,5 x 10^2 CFU/gr of faecal content) and 38% (1.5 × 10^1 CFU/gr of faecal content) of mice, respectively (Figure 2D). Notably, the first evidence of systemic dissemination was observed 3 days post-infection, as 50% of mice showed moderate bacterial load in the liver (5.5 × 10^3 CFU/gr) and 13% also exhibited splenic infection. Seven days after infection, a marked increase in bacterial loads was registered in distant organs with 84% of mice showing viable counts reaching 5.5 x 10^4 CFU/ml in the blood (P ≤ 0.01 day 1 vs. day 7 and day 3 vs day 7; Figure 2C) and 1.2 x 10^7 CFU/gr in liver (P ≤ 0.001 day 1 vs. day 7 and P ≤ 0.05 day 3 vs day 7; Figure 2C). Furthermore, all mice showed sustained bacterial presence in the spleen, with mean values of 3.9 x 10^6 CFU/organ (P ≤ 0.001 day 1 vs. day 7 and day 3 vs day 7; Figure 2C). At this time point, all infected mice exhibited the highest bacterial burden in intestinal compartments, with mean values of 5.4 x 10^7 CFU/gr of faecal content in the cecum (P ≤ 0.05 day 1 vs. day 7 and P ≤ 0.01 day 3 vs day 7; Figure 2D) and 7.9 x 10^6 CFU/gr of faecal content in the colon (P ≤ 0.01 day 1 vs. day 7 and day 3 vs day 7; Figure 2D).

**Figure 2:**
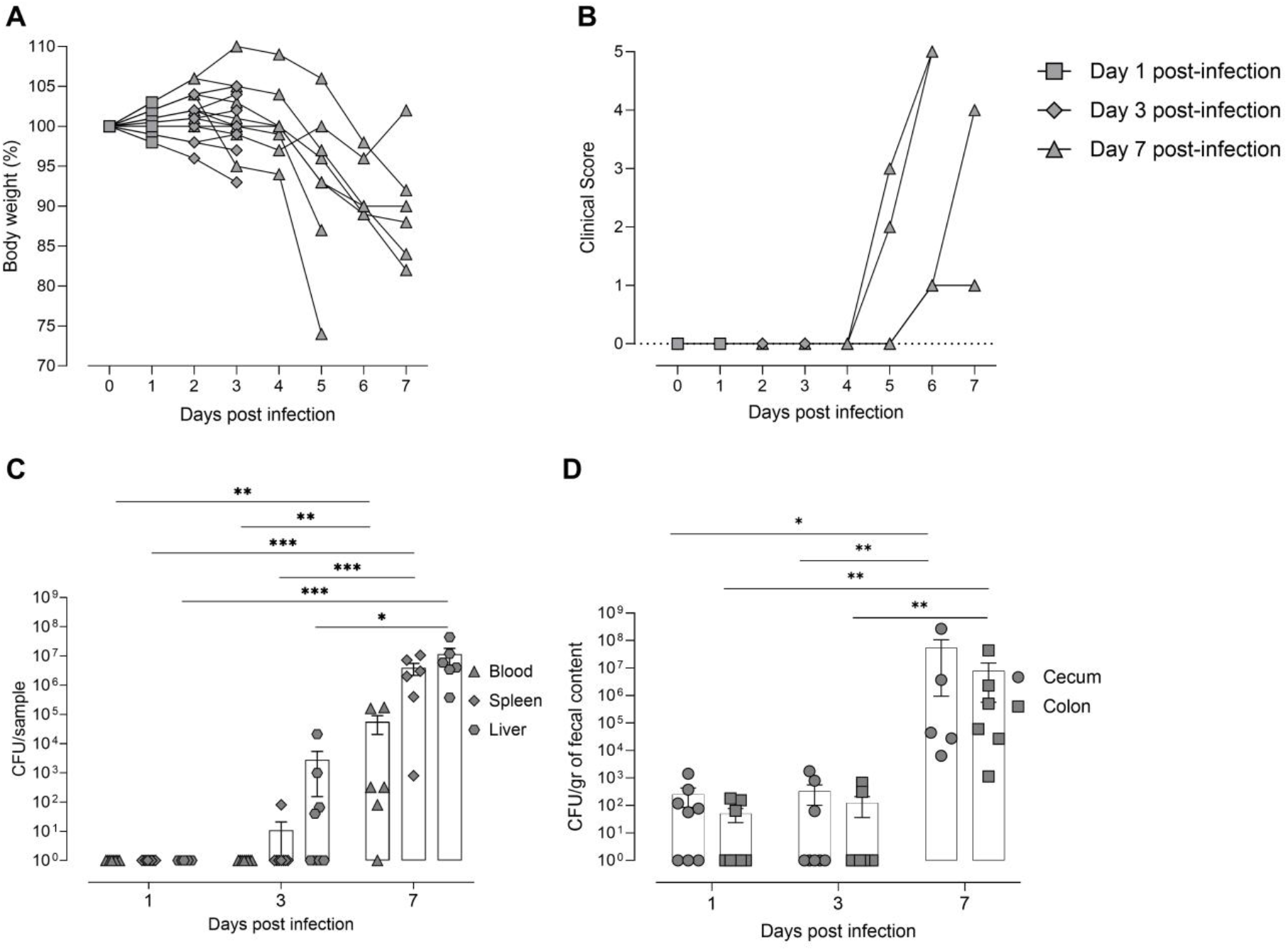
Time-course analysis of body weight, clinical scores and bacterial dissemination in mice infected with *Salmonella* Typhimurium D23580. Animals from Cohort 1A were intragastrically infected by oral gavage with 10^8 CFU/mouse of *S.* Typhimurium D23580 and sacrificed at 1, 3 and 7 days after infection. **(A-B)** Mice were daily monitored until sacrifice for changes in body weight **(A)** and clinical scores **(B)**. Body weight loss was expressed as the percentage of body weight change in comparison to the weight prior to infection, and clinical scores were expressed in a scale ranging from 0 (normal) to 5 (clinical signs requiring euthanasia according to the predefined humane endpoint criteria). Symbols represent individual mice. In panels A and B, squares indicate mice assigned to the day 1 sacrifice group, diamonds indicate mice assigned to the day 3 sacrifice group, and triangles indicate mice assigned to the day 7 sacrifice group. **(C-D)** Bacterial loads in blood, spleen, liver **(C)**, and cecum and colon **(D)** are shown. Symbols represent individual mice, and bars indicate the mean value ± SEM of CFU/ml of blood, CFU/gr of liver/spleen, and CFU/gr of faecal content in cecum and colon. Statistical analysis was performed using the Kruskal–Wallis test followed by Dunn’s multiple-comparisons test. *P ≤ 0.05, **P ≤ 0.01, ***P ≤ 0.001.

### *S*. Typhimurium D23580 infection induces progressive intestinal and systemic tissue damage

Histopathological analysis of cecum, liver, and spleen were performed in Cohort 1A to evaluate tissue damage at 1, 3, and 7 days post-infection following infection with D23580. An additional group of four naïve, uninfected mice served as the day-0 reference control group. Consistent with the original scoring system, minimal cecal alterations detected in naïve, uninfected controls were interpreted as low-grade baseline changes and were not considered indicative of disease, as previously reported (23). In the cecum, a significant increase in the total histopathological score was observed at 3 and 7 days post-infection compared to the naïve group (P ≤ 0.05; Figure 3F). No significant differences were observed among the infected groups at the evaluated time-points. Inflammation, reflecting inflammatory cell infiltration, significantly increased at 3 and 7 days post-infection compared to day 0 (P ≤ 0.01 and P ≤ 0.05, respectively), whereas submucosal edema and goblet cell depletion augmented over time without reaching statistical significance. Epithelial integrity showed no significant variation across infected groups over time (Figure 3F).

**Figure 3:**
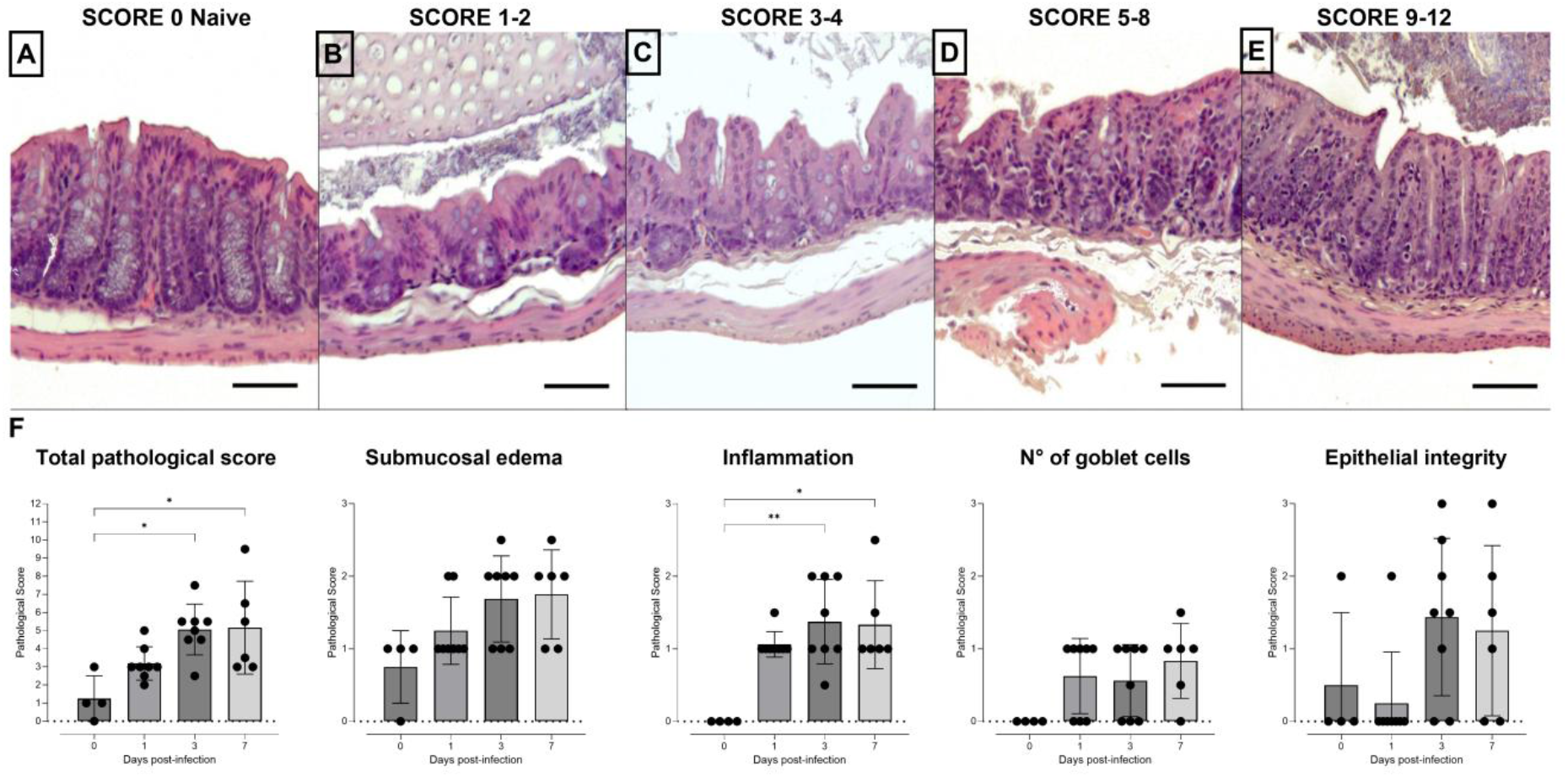
Histopathological scoring of cecal lesions. **(A-E)** Representative hematoxylin and eosin-stained sections of cecum (magnification 200×, scale bar: 100 µm) collected from Cohort 1A and naïve control mice. Histopathological scoring (from 0 to 12) was based on semiquantitative assessment of 4 parameters (submucosal edema, inflammatory infiltrate, goblet cell depletion, and epithelial integrity; Supplementary Table S1). Five representative panels illustrate increasing histopathological damage: **(A)** score 0 (no alterations, naïve mice); **(B)** scores 1–2 (minimal damage); **(C)** scores 3–4 (mild damage); **(D)** scores 5–8 (moderate damage); and **(E)** scores 9–12 (severe damage); **(F)** Total (0-12) and individual (0-3) histopathological scores of the 4 parameters analyzed at 1, 3, and 7 days post-infection, compared to naïve controls. Each dot represents a single mouse; bars indicate mean ± SEM. Kruskal-Wallis test, followed by Dunn’s test for multiple comparisons, was used for assessing statistical differences amongst groups. *P ≤ 0.05; **P ≤ 0.01.

In liver and spleen, histopathological scoring revealed minimal alterations at days 1 and 3 post-infection, whereas marked inflammatory and necrotic lesions were observed at day 7 (Figure 4). At this time point, histopathological scores were significantly increased compared with naïve controls (P ≤ 0.01). Damage was also significantly higher on day 7 compared with both days 1 and 3 post-infection in both organs (P ≤ 0.01 and P ≤ 0.001) (Figure 4I-J). Although no statistically significant differences were detected compared to the naïve group, vascular thrombosis was observed as an additional finding in both liver and spleen at 7 days post-infection (data not shown). Specifically, vascular thrombi were detected in the liver of 2/6 mice (Fisher’s exact test, P = 0.4667) and in the spleen of 4/6 mice (Fisher’s exact test, P = 0.0762), compared with naïve controls. No vascular thrombi were detected in either liver or spleen of naïve controls or mice sacrificed at days 1 and 3 post-infection.

**Figure 4.**
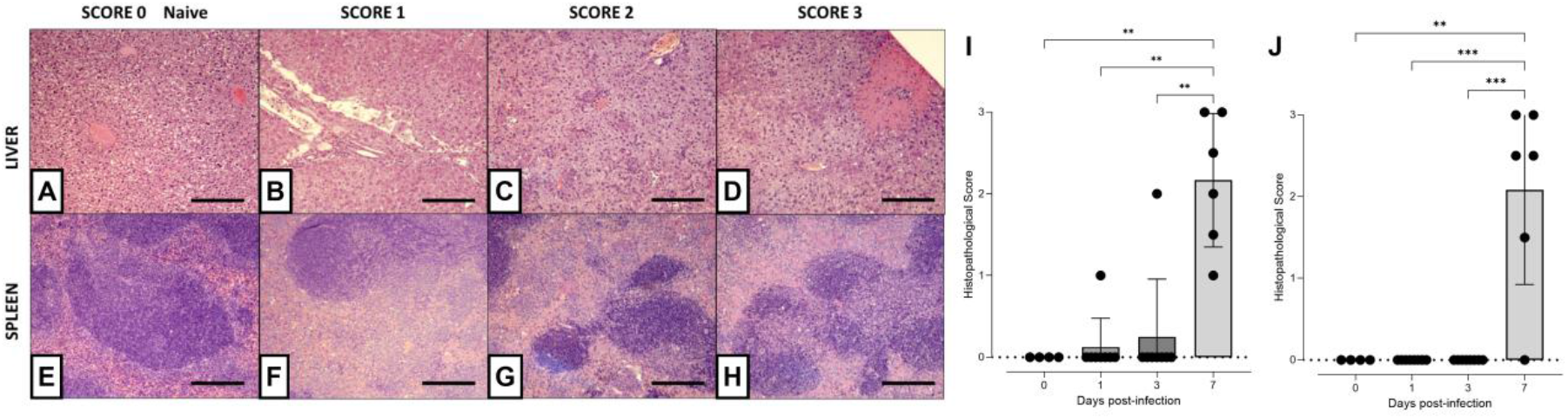
Histopathological scoring of liver and spleen. **(A–H)** Representative haematoxylin and eosin-stained sections illustrating histopathological scoring in liver **(A–D)** and spleen (**E–H)** (magnification 100×, scale bar: 200 µm) collected from Cohort 1A and naïve control mice. Histopathological scoring (from 0 to 3) was based on semiquantitative assessment of inflammation and necrosis in liver and spleen, according to organ-specific criteria (Supplementary Table S2). Liver scores ranged from 0 (normal architecture) to 3 (severe inflammation and necrosis), reflecting increasing hepatocyte necrosis and infiltration by histiocytes, neutrophils, and macrophages. Spleen scores ranged from 0 (naïve) to 3, the latter characterized by progressive infiltration of the marginal zone and red pulp by macrophages and loss of follicular organization **(A-H)**. **(I–J)** Semiquantitative histopathological scores (0-3) in liver **(I)** and spleen **(J)**, based on the assessment of inflammation and necrosis according to organ-specific criteria (Supplementary Table S2). Dots represent individual mice; bars indicate mean ± standard deviation (SD). Statistical differences between groups were assessed by Kruskal–Wallis test, followed by Dunn’s test for multiple comparisons. **P ≤ 0.01; ***P ≤ 0.001.

### KC and TNF-α expression increase at later stages of infection

The local inflammatory response induced by *S*. Typhimurium was characterized in mice from Cohort 1A by qRT-PCR analysis of KC and TNF-α mRNA expression levels in the cecum and liver at 1, 3 and 7 days post-infection, using naïve animals as the reference group (day 0) (Figure 5). KC, the murine functional homologue of human IL-8 (46), and TNF-α (47), a key pro-inflammatory cytokine both showed a significant increase over time. In the cecum, KC expression was significantly higher at day 7 than at days 0 (P ≤ 0.001), 1 (P ≤ 0.001) and 3 (P ≤ 0.01), while TNF-α expression was significantly higher at day 7 than at day 1 (P ≤ 0.05) (Figure 5A-B). In the liver, both KC and TNF-α expression were significantly increased at day 7 compared with all earlier time points (P ≤ 0.05 to P ≤ 0.001) (Figure 5C-D).

**Figure 5.**
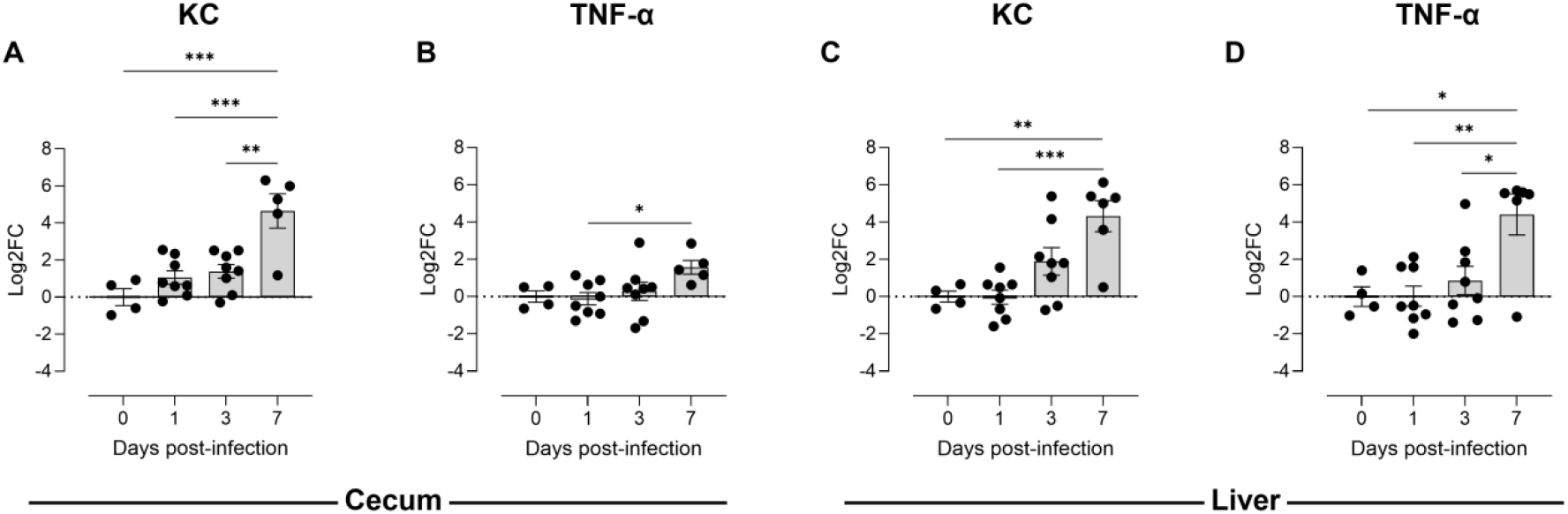
Expression of KC and TNF-α genes in cecum and liver of *S.* Typhimurium D23580-infected mice. The relative mRNA expression levels of KC and TNF-α were quantified in the cecum **(A, B)** and liver **(C, D)** of mice from Cohort 1A. Gene expression was normalized to GAPDH and expressed as log2 fold change (Log2 FC) relative to naïve animals. Data are shown as mean ± SEM. Statistical analysis was performed using one-way ANOVA followed by Tukey’s post hoc test for multiple comparisons. *P ≤ 0.05, **P ≤ 0.01, ***P ≤ 0.001.

### Serum cytokine/chemokine profiling supports delayed systemic inflammatory activation

Serum cytokine and chemokine kinetics were characterized by integrating data from Cohorts 1A and 1B. Cohort 1A provided measurements at days 1, 3 and 7 post-infection, whereas Cohort 1B provided additional measurements at days 3, 5 and 7, allowing characterization of serum mediator dynamics from day 0 (pre-infection serum samples) to day 7 post-infection (Figure 6A).

**Figure 6.**
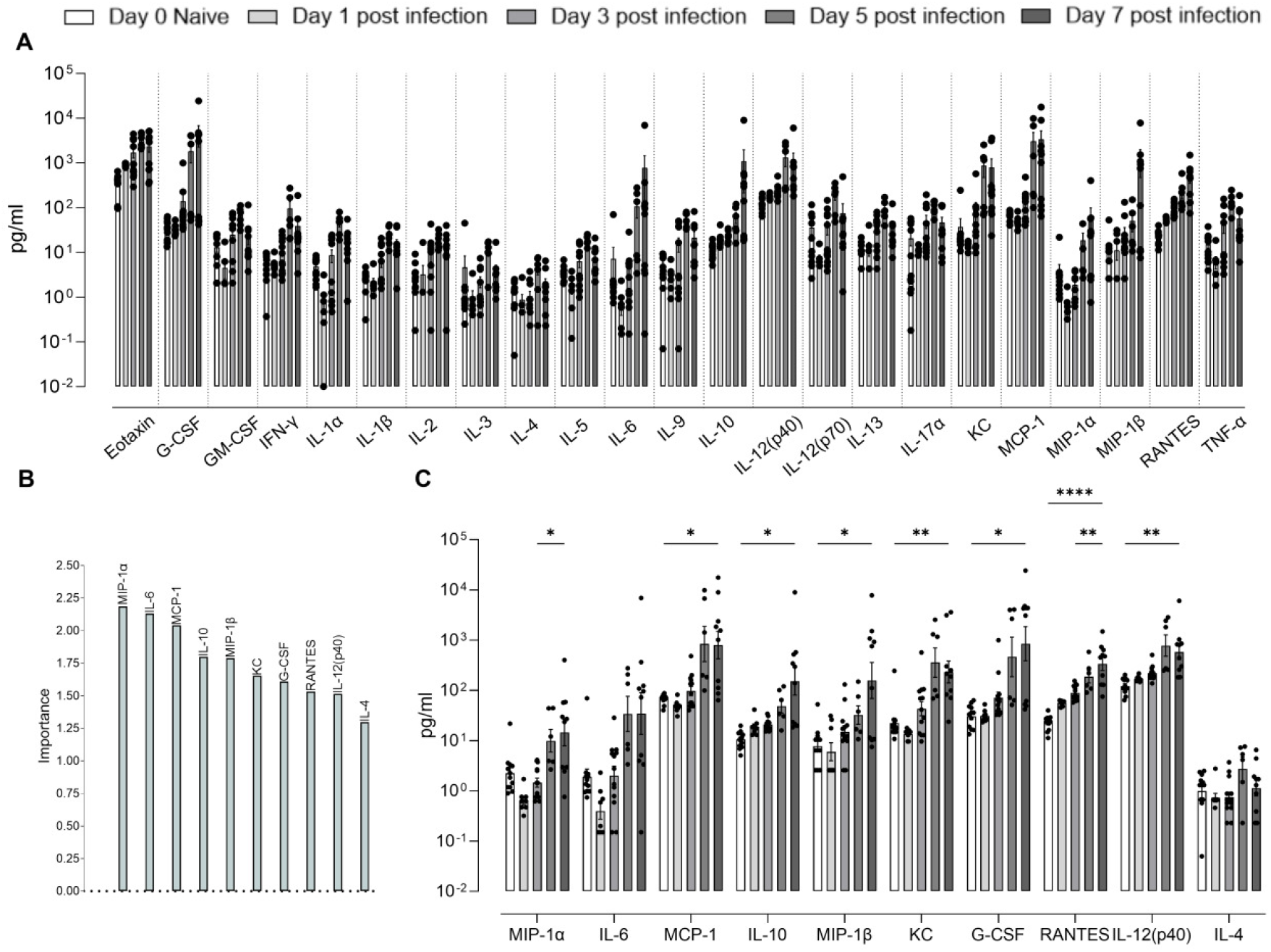
Serum cytokine/chemokine dynamics and machine learning-based prioritization of mediators associated with surrogate mortality-risk labels. Serum concentrations of 23 cytokines/chemokines were measured in samples from Cohorts 1A and 1B using a 23-plex Luminex assay. **(A)** Concentrations of all 23 analytes from day 0 to day 7 post-infection. Day 0 represents the pre-infection, naïve reference group. Data are shown as mean ± SEM, with individual animals represented by dots. **(B)** Random Forest variable-importance scores for the ten highest-ranked mediators in the multivariate classification of the surrogate mortality-risk labels. **(C)** Temporal profiles of the ten highest-ranked mediators. Concentrations are expressed as pg/mL and were calculated from analyte-specific standard curves. For temporal comparisons, mediator concentrations were log10-transformed and analysed separately using ordinary one-way ANOVA followed by Tukey’s multiple-comparisons test. Complete pairwise comparisons for all 23 analytes are reported in Supplementary Table S3. The complete machine-learning workflow and model-performance metrics are reported in the Supplementary Methods and Supplementary Results.

Across the 23 analytes, the most pronounced changes were generally observed at later stages of infection. Chemokines involved in leukocyte recruitment, including KC, MCP-1, MIP-1α, MIP-1β and RANTES, showed significant increases compared with baseline, particularly at days 5 and/or 7 (Supplementary Table S3). Among these, RANTES displayed one of the most sustained responses, with significantly higher concentrations than baseline at all post-infection time points, whereas Eotaxin showed an earlier increase relative to baseline. IL-6, G-CSF and IL-12(p40) also showed consistent late changes. TNF-α, IFN-γ, IL-1β and IL-12(p70) displayed their most pronounced differences in comparisons involving day 5. IL-1α showed an early increase at day 1 followed by additional differences at later time points, while IL-13 rose sharply at day 5 alone, returning to baseline by day 7. IL-10 showed a predominantly late increase, particularly at day 7, whereas IL-4 did not show significant differences across the analysed time points.

Overall, these data indicate a heterogeneous systemic response involving inflammatory cytokines, leukocyte-recruiting chemokines and immunoregulatory mediators. The delayed increase in several circulating mediators was consistent with the progression of bacterial dissemination, tissue pathology and local KC and TNF-α expression. Complete pairwise comparisons for all 23 analytes are reported in Supplementary Table S3.

To further characterize the systemic host response during the *Salmonella* infection, as an exploratory analysis, we examined whether cytokine and chemokines profiles were associated with mortality risk. Since mortality could not be measured directly in the study, we first derived a surrogate mortality label for each animal based on results obtained in a previous study (as detailed in Supplementary Methods and Results) (42). Once the surrogate mortality-risk labels were computed, four new classifiers (Random Forest, Support Vector Machines, XGBoost and logit) were trained using the cytokine/chemokine levels as predictors and the surrogate labels as outcome. Among the four classifiers tested, the best-performing model was a random forest, which was therefore used to summarize the relative contributions of individual mediators to the multivariate classification.

Variable importance analysis identified a core panel of 10 different cytokines/chemokines as major contributors to model decision (Figure 6B). Most of these mediators also displayed statistically significant temporal changes, with marked increases at later stages of infection (Figure 6C; Supplementary Table S3). The prioritized panel included both pro-inflammatory markers (MIP-1α, MIP-1β, IL-6, MCP-1, KC, RANTES) and regulatory mediators (IL-10, IL-4, G-CSF, IL-12(p40)). Thus, the model prioritized a mixed inflammatory and regulatory serum profile rather than a set restricted to canonical pro-inflammatory mediators. The corresponding univariate comparisons between the two surrogate mortality-risk label groups and the performance metrics of the evaluated classifiers are reported in Supplementary Tables S4 and S5, respectively.

### Inactivated *S*. Typhimurium D23580 induces robust antigen-specific IgG and memory B-cell responses

To demonstrate the applicability of the present model for evaluating vaccine-induced immunity against iNTS infection, serum samples from vaccinated mice in Cohorts 1C and 2 were analysed. Following the immunization with an inactivated *S.* Typhimurium D23580 prototype vaccine, serum anti-*S.* Typhimurium IgG responses were assessed.

As depicted in Figure 7A, anti-*S.* Typhimurium IgG levels increased progressively following immunization (Figure 7A), with a significant rise already detectable at week 3 (GMT = 3,098; 95% CI 2,086–4,599; P ≤ 0.001 vs week 0), followed by a further increase at week 6 (GMT = 44,871; 95% CI 32,630–61,705; P ≤ 0.001 vs week 3 and P ≤ 0.0001 vs week 0), and reaching peak levels at week 9 (GMT = 155,540; 95% CI 116,888–206,974; P ≤ 0.05 vs week 6; P ≤ 0.0001 vs weeks 3 and 0).

**Figure 7.**
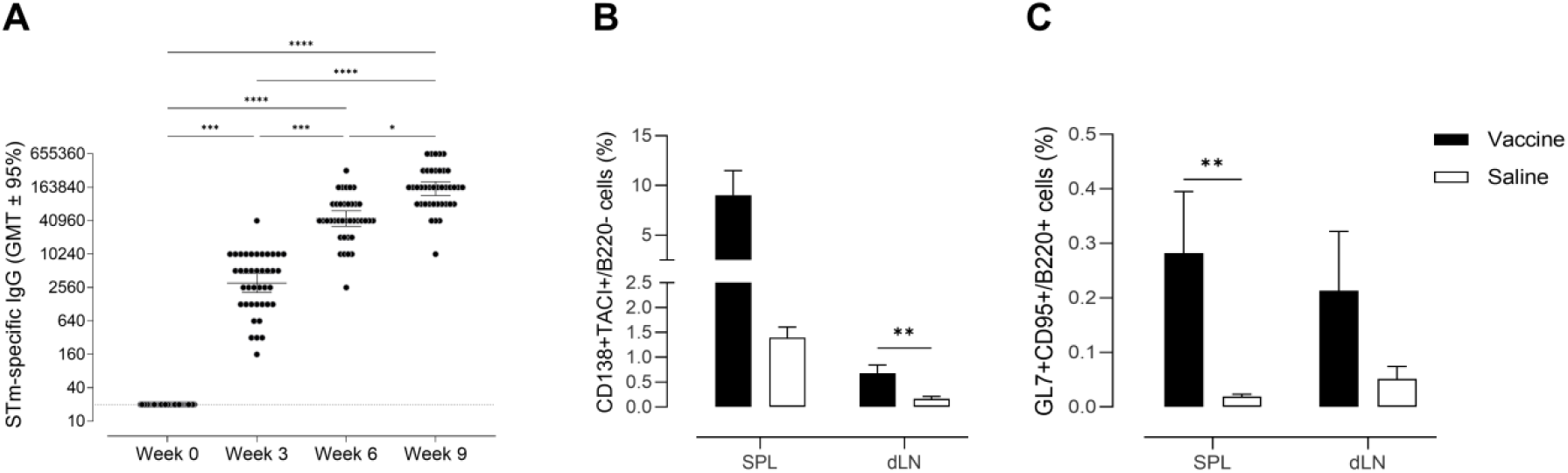
Vaccine-induced serum IgG responses and memory B-cells. C57BL/6 mice were immunized subcutaneously at weeks 0, 3 and 6 with formalin-inactivated *Salmonella* Typhimurium D23580. **(A)** Time course of *S.* Typhimurium-specific serum IgG responses, measured by ELISA at weeks 0 (pre-immunization), 3, 6 and 9 using serum samples from vaccinated mice in Cohorts 1C and 2. Antibody titers are expressed as the reciprocal of the highest serum dilution yielding an optical density at least twofold higher than background. Data are presented as geometric mean titers (GMT) with 95% confidence intervals (CI), and symbols represent individual mice. The dashed horizontal line indicates the lower limit of detection (LLOD = 20). **(B–C)** Flow-cytometric characterization of selected B-cell populations in Cohort 2. Six months after the first immunization, vaccinated (n = 8) and saline-treated control mice (n = 6) were challenged intragastrically with 10^8 CFU/mouse of live *S.* Typhimurium D23580 and euthanized six days later. Frequencies of IgD⁻B220⁻CD138⁺TACI⁺ plasma cells **(B)** and B220⁺GL7⁺CD95⁺ germinal-center B cells **(C)** were determined in the spleen and draining lymph nodes by multiparameter flow cytometry. Bars represent mean ± SEM. Statistical differences in serum IgG titers **(A)** were assessed using the Kruskal-Wallis test, followed by Dunn’s post-test for multiple comparisons were used. *P ≤ 0.05; **P ≤ 0.01; ***P ≤ 0.001; ****P ≤ 0.0001, whereas differences between vaccinated and saline-treated mice in the flow-cytometry analyses **(B–C)** were assessed using exact two-tailed Mann–Whitney U tests. *P ≤ 0.05; **P ≤ 0.01; ***P ≤ 0.001; ****P ≤ 0.0001.

The murine model also proved suitable for the long-term characterization of vaccine-induced cellular immune responses and for investigating their rapid reactivation upon subsequent pathogen exposure. To this end, mice were immunized subcutaneously at weeks 0, 3 and 6 with formalin-inactivated *S*. Typhimurium D23580, while saline-treated mice served as controls. Six months after the first immunization, vaccinated and control mice were challenged by oral gavage with live *S*. Typhimurium D23580 and euthanized 6 days later. Selected B-cell populations were then characterized in the spleen and draining lymph nodes by multiparameter flow cytometry. The frequency of IgD⁻B220⁻CD138⁺TACI⁺ plasma cells was higher in vaccinated mice, reaching statistical significance in the draining lymph nodes (P ≤ 0.01; Figure 7B). Germinal-center B cells, defined as B220⁺GL7⁺CD95⁺ cells, were also increased in vaccinated mice, with a statistically significant difference observed in the spleen (P ≤ 0.01; Figure 7C). Overall, vaccination with formalin-inactivated D23580 was associated with differences in B-cell populations after challenge, characterized by increased plasma-cell frequencies in the draining lymph nodes and increased germinal-center B-cell frequencies in the spleen.

### Vaccination reduces disease severity and bacterial dissemination after intragastric challenge

To evaluate vaccine-associated protection against iNTS disease, a challenge subset of Cohort 1C consisting of vaccinated mice (n = 8) and saline-treated control mice (n = 8) was intragastrically challenged by oral gavage with 10^8 CFU/mouse of live *S*. Typhimurium D23580 and monitored for clinical outcomes (Figure 8). As shown in Figure 8A, vaccinated mice maintained stable body weight over time, whereas control animals showed a progressive weight loss starting from day 3 post- infection (Figure 8A). Consistently, clinical scores remained low in vaccinated mice, with only two animals showing a slightly higher clinical score (score = 1) at day 6 post-infection, while control mice developed severe disease (scores up to 4–5), leading to the death of two animals by day 6 (Figure 8B). To further evaluate vaccine protection, bacterial dissemination from the infection site was quantified at day 6 post-infection in both intestinal and extra-intestinal compartments (Figure 8C–D). Vaccinated mice showed significantly lower bacterial loads in blood and liver compared to controls (Figure 8C). Bacteraemia was detected in 25% of vaccinated animals, compared to 70% of control mice, and bacterial loads were also lower in vaccinated than in saline-treated animals (10^3 vs. 10^5 CFU/mL; P ≤ 0.05). Similarly, multiplication in the liver was reduced in vaccinated animals (10^5 vs. 10^7 CFU/gr; P ≤ 0.05). A marked decrease of bacterial dissemination was also observed in intestinal compartments of vaccinated mice compared to controls, with a statistically significant CFU reduction in the colon of vaccinated (10^4 CFU/gr faecal content) compared to control (10^8 CFU/gr) mice (P ≤ 0.05), (Figure 8D). Interestingly, bacterial shedding in the faeces was observed in only 50% of vaccinated mice (up to 10^5 CFU/gr) and 85% of saline-injected mice (up to 10^9 CFU/gr). Overall, these findings support the suitability of the present model for evaluating vaccine-induced protection against systemic *Salmonella* infection, as shown by measurable vaccine-associated effects, including reduced bacterial dissemination and attenuation of clinical disease after challenge.

**Figure 8:**
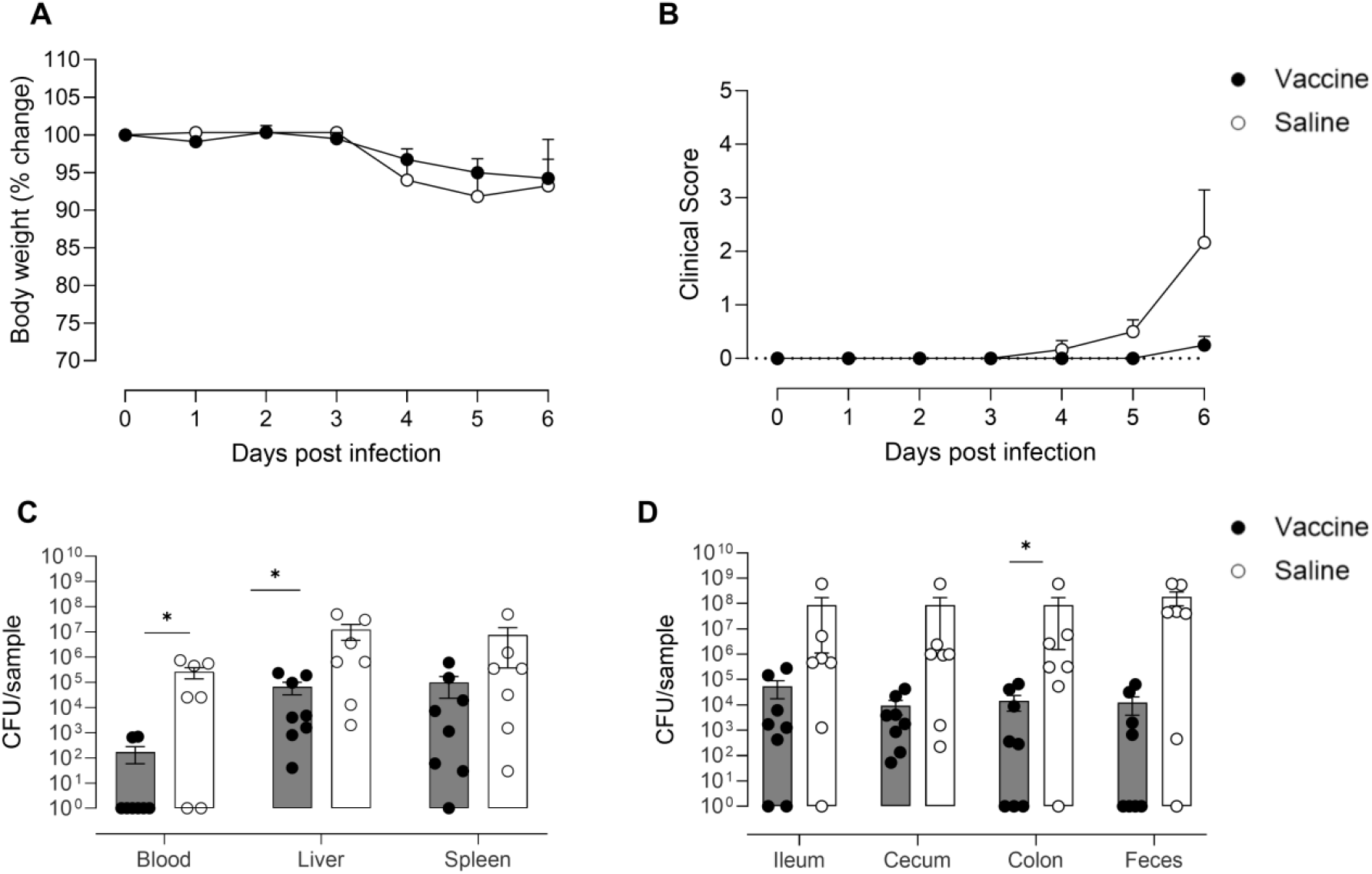
Impact of vaccination with inactivated *S*. Typhimurium on clinical signs and bacterial dissemination. Mice from Cohort 1C were subcutaneously immunized with 10^8 CFU equivalent/mouse of formalin-inactivated *S*. Typhimurium D23580 at weeks 0, 3 and 6, while control animals received saline. Ten weeks after initial immunization, a challenge subset of Cohort 1C comprising both vaccinated (n = 8; solid symbols) and control mice (n = 8; open symbols) was intragastrically infected by oral gavage with 10^8 CFU/mouse of live D23580 bacteria and sacrificed 6 days post-infection. **(A)** Body weight loss is expressed as the percentage of the body weight change in comparison with the weight prior to infection and reported as group mean ± SEM. **(B)** Mice were daily monitored, and clinical scores expressed in a scale ranging from 0 (normal) to 5 (clinical signs requiring euthanasia according to the predefined humane endpoint criteria) and reported as group mean ± SEM. **(C)** Bacterial viable counts are reported as the number of CFU/ml of blood, CFU/liver, CFU/spleen. **(D)** Viable counts are reported as CFU/gr of faecal content in ileum, cecum and colon, and CFU/gr of feces. Symbols represent individual mice: solid symbols for vaccinated and open for controls, while bars indicate the mean value ± SEM. Statistical analysis was performed using the two-tailed Mann–Whitney U test for multiple pairwise comparison. *P ≤ 0.05.

## DISCUSSION

In the present study, an intragastric model of invasive non-typhoidal *Salmonella* (iNTS) disease was characterized using susceptible Nramp1-/-C57BL/6 mice infected with the ST313 clinical isolate *Salmonella enterica* serovar Typhimurium D23580. By administering the bacteria intragastrically without antibiotic pretreatment or gastric acid neutralization, this model retains the gastrointestinal phase of infection, allowing D23580 to reach and colonize the intestine before disseminating to extraintestinal sites. It thereby provides an integrated view of iNTS infection dynamics, capturing the transition from intestinal infection to systemic dissemination and subsequent tissue injury at sites of *Salmonella* replication, while avoiding the direct bypass of the intestinal compartment inherent to parenteral challenge models and the deliberate disruption of microbiota-associated colonization resistance typical of antibiotic-treated models (22, 23, 48, 49). These features support its use as a preclinical tool for investigating iNTS disease progression and evaluating vaccine-induced immune responses.

Following intragastric challenge, *S.* Typhimurium D23580 exhibited a progressive dissemination pattern characterized by an initial phase of localised intestinal infection followed by systemic spread. Bacteria were detected in the cecum and colon during the early phase of infection, whereas liver involvement became evident by day 3. By day 7, most animals showed substantial bacterial burdens in the blood, liver and spleen, together with increased intestinal loads and progressive clinical deterioration. The simultaneous persistence of bacteria in the intestine and their accumulation in systemic organs is consistent with the typhoid-like behaviour previously described for the D23580 strain (33). Early dissemination may occur through the uptake of *Salmonella* at specialized intestinal sites, including Peyer’s patches and M cells, or by CD18+ phagocytic cells, thus facilitating its entry into underlying tissues and lymphoid compartments (50).

The dissemination kinetics observed here differed from those reported in streptomycin-pretreated mouse models, in which antibiotic-mediated disruption of the intestinal microbiota promotes rapid intestinal colonization, pronounced inflammation and early systemic spread (22, 23, 49). The slower progression observed in the present model may reflect, at least in part, the absence of antibiotic-mediated disruption of microbiota-associated colonization resistance. Recent work in antibiotic-naïve mice has shown that *S.* Typhimurium can nevertheless overcome colonization resistance through infection-induced changes in the intestinal environment, including increased epithelial oxygenation and altered luminal metabolite availability, before measurable changes in microbiota composition become apparent (51). Although these parameters were not assessed in the present study, these findings provide a possible biological context for infection establishment and progression in the absence of antibiotic pretreatment. Overall, the model offers a gradual intestinal-to-systemic disease trajectory that may be useful for investigating typhoid-like iNTS pathogenesis under conditions that do not involve deliberate antibiotic disruption of the intestinal environment.

The progressive systemic bacterial dissemination was associated with increasingly severe histopathological alterations. Histological analysis demonstrated a gradual escalation of inflammatory lesions across both gastrointestinal and extraintestinal compartments. In the cecum, the total histopathological score was significantly increased at days 3 and 7 post-infection compared with naïve controls, largely reflecting increased inflammatory infiltration. In contrast, hepatic and splenic lesions were limited during the early stages of infection and became more pronounced at day 7, when inflammatory and necrotic changes were significantly increased. Vascular thrombi were also detected in a subset of animals at this late time point, consistent with previous reports of *Salmonella*-associated thrombosis in mice (38), although the differences compared with naïve controls were not statistically significant. Overall, histopathological alterations closely paralleled bacterial loads and clinical deterioration over time, supporting an association between *Salmonella* replication, inflammation and tissue injury during disease progression.

To further investigate the molecular correlates of the histopathological changes, the expression of key mediators of innate immune activation was examined in the cecum and liver. Both KC, the murine functional counterpart of human IL-8 and a major neutrophil chemoattractant (46, 52, 53), and TNF-α, a central mediator of inflammatory-cell recruitment and macrophage activation (47, 54), displayed a progressive but delayed induction compared with the inflammatory responses described in streptomycin-pretreated models (53, 55). In particular, no significant upregulation of KC was observed during the early stages of infection, despite intestinal colonization and initial dissemination to the liver. This delayed inflammatory response is consistent with the relatively low bacterial burdens at early time points and with a slower engagement of host inflammatory pathways in the absence of microbiota disruption. By day 7 post-infection, increased KC expression in both the cecum and liver, together with increased hepatic TNF-α expression, occurred in the context of high systemic bacterial burdens and marked tissue pathology (53–55). Collectively, these findings suggest that tissue injury emerged progressively alongside increasing bacterial replication and intensification of local inflammatory responses during systemic infection. The distinct but coordinated responses observed in the cecum and liver are consistent with compartment-specific contributions to disease progression, with the late hepatic response coinciding with systemic bacterial dissemination and marked tissue pathology.

Serum cytokine and chemokine profiling further supported a delayed and heterogeneous systemic immune response during D23580 infection. The most consistent changes occurred 7 days after infection and involved chemokines associated with leukocyte recruitment, including KC, MCP-1, MIP-1α, MIP-1β and RANTES, together with IL-6, G-CSF and IL-12p40. Eotaxin showed an earlier increase relative to baseline, whereas TNF-α, IFN-γ, IL-1β and IL-12p70 displayed their most pronounced differences in comparisons involving day 5. IL-10 increased predominantly at later time points, particularly at day 7, while IL-4 did not show significant changes. Overall, the serum profile was not consistent with a uniform pro-inflammatory cytokine burst, but rather with the coordinated modulation of inflammatory, leukocyte-recruiting and immunoregulatory mediators (53, 55–60). The temporal behaviour of TNF-α was particularly notable: serum concentrations peaked at day 5 and were numerically lower at day 7, although this difference was not statistically significant. This pattern does not necessarily indicate resolution of inflammation, as circulating concentrations may not reflect the persistence of TNF-α activity within infected tissues, particularly when local inflammatory gene expression and tissue pathology remain evident (54). The concomitant late increase in IL-10 may reflect the induction of counter-regulatory mechanisms during progressive infection; however, IL-10 can also promote systemic *S.* Typhimurium infection by attenuating antimicrobial immune responses (61). The absence of significant changes in IL-4 suggests that this regulatory response was not accompanied by a generalized increase in type 2-associated cytokines (60). This coordinated inflammatory and regulatory profile is broadly consistent with observations in children with acute invasive non-typhoidal *Salmonella* disease. In Malawian children, severe iNTS was associated with cytokine patterns combining macrophage-related, inflammatory and regulatory responses, while fatal disease was associated with increased IL-8 and HGF and altered IL-1 signalling (62). Although murine and human cytokine concentrations cannot be compared directly, the increase in KC provides a relevant functional point of comparison with the IL-8-associated neutrophil-recruitment response described in human iNTS.

To validate the present infection model as a tool for preclinical evaluation of novel vaccine candidates, a prototype formalin-inactivated *S.* Typhimurium D23580 vaccine was prepared and its immunogenicity and protective efficacy were assessed.

Vaccination elicited robust *Salmonella-*specific IgG responses, with antibody titers progressively increasing over time, indicating effective induction of humoral immunity. Although cellular immunity is considered essential for protection against intracellular *Salmonella*, antibodies may contribute to bacterial control by promoting complement-mediated killing and limiting bacterial dissemination (32, 63–66).

Following intragastric challenge with live D23580, vaccinated animals showed a delayed disease onset, reduced clinical severity and lower bacterial burdens in both extraintestinal and intestinal compartments than saline-treated controls. Particularly notable was the reduction in intestinal bacterial loads and faecal shedding in vaccinated animals. Because intestinal persistence and faecal shedding are relevant features of enteric *Salmonella* infection (67), these findings suggest that vaccination may limit not only systemic dissemination but also the intestinal bacterial reservoir, with the potential to reduce faecal-oral transmission (3, 4). Previous vaccination studies have frequently relied on resistant mouse strains or antibiotic pretreatment to enhance intestinal infection and improve experimental reproducibility. Although these approaches facilitate onset of infection, they may also alter host-pathogen interactions and influence the evaluation of vaccine efficacy (33). In contrast, vaccine-associated protection was demonstrated here using an intragastric challenge model that preserves the intestinal microbiota and the natural route of infection, thereby more closely mimicking the iNTS disease in humans.

To study long-term vaccine-induced B-cell responses, and to investigate the rapid reactivation upon pathogen exposure, selected B-cell populations were analysed six days after D23580 challenge administered six months after the first immunization. At this time point, vaccinated mice showed significantly higher frequencies of IgD⁻B220⁻CD138⁺TACI⁺ plasma cells in the draining lymph nodes and of B220⁺GL7⁺CD95⁺ germinal-center B cells in the spleen compared with saline-treated controls. Since both groups received the same challenge, these differences suggest that previous immunization influenced the B-cell response observed after infection.

The higher frequencies of these B-cell populations in vaccinated mice are consistent with a secondary response following pathogen re-exposure. Previous studies have shown that antigen re-exposure can induce early plasma-cell differentiation and germinal-center responses, supporting the possibility of a secondary B-cell response following vaccination (68). Cunningham *et al.* showed that germinal-center formation following primary infection of naïve mice with attenuated *S.* Typhimurium was delayed until several weeks after infection (66), further supporting this interpretation, although in a different model. However, *S.* Typhimurium infection can also disrupt established germinal-center responses (69), and differences in the effects of infection on B-cell populations may have contributed to the findings observed here.

Some limitations should be considered in the present study. The model was established using a single mouse genetic background, C57BL/6 mice, and a single ST313 clinical isolate, which may limit the generalizability of the findings to other host genotypes and *Salmonella* lineages. Although the infection was performed without antibiotic pretreatment, the composition and function of the intestinal microbiota were not directly characterized. The challenge study of vaccinated mice was conducted in relatively small cohorts.

In conclusion, the murine model of *S.* Typhimurium D23580 infection described here recapitulates key features of the clinical progression and pathophysiology of invasive iNTS disease, including the transition from intestinal infection to systemic dissemination, tissue injury and delayed inflammatory activation (32, 70). By avoiding antibiotic pretreatment, the model enables a gradual course of infection without deliberate depletion of the intestinal microbiota, providing a suitable tool for investigating host-pathogen interactions, infection kinetics and the inflammatory and immune responses associated with disease progression. The immune response observed after administration of the prototype formalin-inactivated *Salmonella* vaccine provides a proof-of-concept evidence that the model is a useful tool able to dissect the effects of vaccination in clinical disease, bacterial dissemination and immune responses. Overall, the integration of clinical, microbiological, histopathological and immunological readouts obtained at different time points after infection, together with exploratory multivariate analyses, supports the use of this model for evaluating potential therapeutic and prophylactic interventions against invasive non-typhoidal *Salmonella*.

## Acknowledgments

This study was supported by the Department of Medical Biotechnologies of the University of Siena (D.M.) and by the PRIN 2022 PNRR project code P2022PJYXR_001 entitled “Investigation of the role of humoral immunity induced by glycoconjugate vaccines in the control of invasive non-typhoidal *Salmonella* infections” (E.P.).

We would like to thank Dr Chiara Coppola and Dr Simone Costagli for their valuable technical support during the animal experiments.

EP, VM, FF, GP, SR and DM conceptualized the study; VM, FF, EP, MT, GD, AL and LM processed the samples and carried out the immunological, molecular and histopathological analysis; GM, VM and EP analysed the data; EP, SP, GP and DM supervised the study; VM, GM, EP, MT and GD wrote the original draft. All the authors read and approved the final version of the manuscript. DM and EP provided financial support.

## Data and code availability

All data, scripts and output files will be made available upon reasonable request to the corresponding author.

